# Mechanical-control of cell proliferation increases resistance to chemotherapeutic agents

**DOI:** 10.1101/2020.01.18.910554

**Authors:** Ilaria Rizzuti, Pietro Mascheroni, Silvia Arcucci, Zacchari Ben-Mériem, Audrey Prunet, Catherine Barentin, Charlotte Rivière, Hélène Delanoë-Ayari, Haralampos Hatzikirou, Julie Guillermet-Guibert, Morgan Delarue

## Abstract

While many cellular mechanisms leading to chemotherapeutic resistance have been identified, there is an increasing realization that tumor-stroma interactions also play an important role. In particular, mechanical alterations are inherent to solid cancer progression and profoundly impact cell physiology. Here, we explore the impact of compressive stress on the efficacy of chemotherapeutics in pancreatic cancer spheroids. We find that increased compressive stress leads to decreased drug efficacy. Theoretical modeling and experiments suggest that mechanical stress leads to decreased cell proliferation which in turn reduces the efficacy of chemotherapeutics that target proliferating cells. Our work highlights a mechanical-form of drug resistance, and suggests new strategies for therapy.

---

Mechanical alterations of solid tumors are a hallmark of cancer progression. Among the many occurring modifications, the most representative forms of mechanical alterations in tumors are changes in extracellular matrix rigidity[1] and build up of compressive stress[2]. Compressive stress accumulation can be found in many cancers such as glioblastoma multiform[3] or pancreatic ductal adenocarcinoma (PDAC)[2].

PDAC is one of the deadliest cancers, with extremely poor prognosis when diagnosed and no efficient treatment available besides surgery. PDAC development is characterized by excessive deposition of extracellular material during which strong modifications of the mechanical environment arises. In particular, the deposition of negatively-charged hyaluronic acid leads to electroswelling of extracellular matrix and subsequent compressive stress experienced by tumor cells[4]. The local growth of cancer cells in an elastic environment also leads to build-up of compressive stress through a process known as growth-induced pressure[5, 6]. PDAC tumors are extremely compressed, within the kPa range[7].

*In vitro*, compressive stress can alter cell physiology in multiple ways, from proliferation[8, 9] to migration[10, 11]. *In vivo*, it has recently been shown that compressive stress built-up in PDAC tumors can exceed blood pressure and participate in blood vessel collapse[12]. Most of large vessels (diameter above 10*μ*m) are clamped, leading to poor perfusion. Vessel collapse is associated with drug resistance: classical first-line chemotherapeutics such as gemcitabine are thought to be unable to reach the tumor, which decreases or even prevents the effect of the drug. Intravenous injection of a pegylated-form of hyaluronidase, an enzyme digesting hyaluronic acid, renormalizes blood vessels and, in combination with gemcitabine, increases chemotherapeutic efficacy[12].

The proposed mechanism overcoming this form of resistance is better tumor perfusion through decrease in compressive stress. However, it remains unclear how the combination of hyaluronidase and gemcitabine really works. Indeed, if the vessels are so collapsed that gemcitabine does not penetrate the tumor, hyaluronidase should not have a better chance to reach the tumor. Another potential explanation which does not depend on perfusion is that compressive stress could directly act on tumor cells and decrease the efficacy of gemcitabine. Experimentally, scarce instances of drug resistance stemming from mechanical stress have been observed for cells growing on substrata of different rigidities[13] or under shear stress[14], with no clear mechanism. In this letter, we wished to explore the paradigm of compression-modulation of drug resistance. We will focus on the situation where perfusion is not an issue: We only investigate the impact of mechanical stress on chemotherapeutic resistance, considering the case of a well-perfused genetically homogeneous tumor, *e.g.* no chemical gradients and no mutation-based resistance.

Given the complexity of a tumor, uncoupling the effect of biochemical and mechanical interactions is a daunting challenge. A good *in vitro* candidate is the multicellular spheroid as a mesoscopic tumor model system: three-dimensional cellular aggregates which remarkably mimic the relevant *in vivo* physiological gradients of mitogens, oxygen, or glucose. They have been extensively used as tumor model systems for the study of drug delivery[15]. Although their mechanical properties might differ from those of tumors, for many purposes, spheroids can be viewed as a tumor subunit. Because they do not have any biochemical crosstalk with their environment, spheroids are ideal to evaluate the impact of mechanical stress on tumor growth. We formed spheroids from a pancreatic Kras^G12D^ cell line, representative of pancreatic cancer mutations[16], using a classical agarose cushion protocol[8] (Fig. 1a). Under normal growth conditions, spheroids grew over time in the hundreds of micrometer size range (Fig. 1b). We restricted ourselves to the use of small spheroids (diameter below 400*μ*m) to avoid chemical gradients as much as possible.

**FIG. 1.**
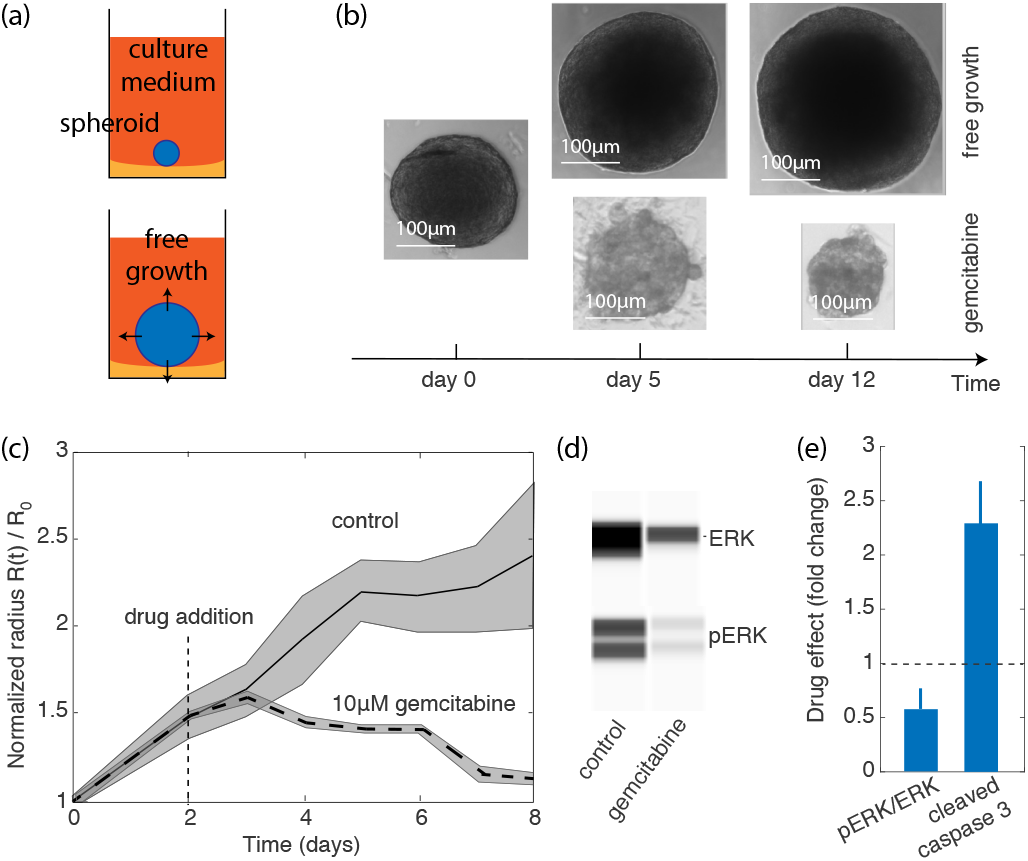
Effect of gemcitabine on the free growth of spheroids. a. Schematic of a free growing spheroid. b. Representative pictures of freely growing pancreatic spheroids and treated with gemcitabine. Drug was added after two days of growth. c. Growth quantification over time. Median values are plotted normalized to time 0, shading corresponding to ± SD over N ≥ 10 spheroids. d. Example of capillary western blot of spheroids after 5 days under drug. e. Analysis of capillary wester blots. Phosphorylated form of ERK (pERK) normalized to total ERK is used as a proxy for the amount of cell proliferation, and cleaved-caspase 3 as a proxy for the amount of dying cells. The data under gemcitabine treatment are normalized to the control. Mean ± SEM over N≥ 3 replicates. All the data were pooled together.

We treated freely growing multicellular spheroids with gemcitabine to investigate the effect of this chemotherapeutic without mechanical stress. Gemcitabine is a cytidine-analog which activates within the cell into a deoxycytidine-triphosphate (dCTP)[17]. dCTP creates, upon DNA incorporation, an irreversible error leading to cell death. Perhaps not surprisingly, we found that spheroids subjected to 10*μ*M gemcitabine decreased in size (Fig. 1b). The decrease was apparent after an average of 1 day post-drug addition (Fig. 1c), consistent with the fact that only S-phase cells are sensitive to the drug. We performed capillary western blots to investigate the changes in proliferating and dying cells under gemcitabine treatment (Fig. 1d, see also Methods for protocol). We measured the ratio between a phosphorylated-form of ERK (pERK) over total ERK as a proxy for cell proliferation[18] and the amount of cleaved-caspase 3 as a proxy for cells undergoing programmed cell death[19] (Fig. 1e). We observed that, as the amount of proliferating cells was halved after 5 days under drug treatment, the amount of dying cells increased by roughly a factor 2, consistent with gemcitabine preferentially killing proliferating cells.

We next sought to investigate the impact of compressive growth-induced pressure on the efficacy of gemcitabine. Several strategies have been developed to study cells under growth-induced pressure, such as microfluidic confining chambers[20], embedding single cells in agarose or PEG-heparin hydrogels[5, 9] or in alginate shells[6], or directly embedding spheroids inside alginate hydrogels[21]. In order to easily follow single spheroids, we opted for a strategy where spheroids were directly embedded into low-melting 1% agarose (see Methods). Briefly, 200*μ*L of a 2% low-melting agarose solution kept at 37°C was mixed with a 200*μ*L solution containing a spheroid, and polymerized on ice to limit the rapid sedimentation of the spheroid which could lead to partial embedding. Gelling on ice was fast and did not affect the spheroid, as assessed by the normal growth of the control subjected to similar conditions.

Measurement of growth-induced pressure requires careful characterization of material property, a point that is often disregarded, as reminded in [9]. Rheological measurements of agarose hydrogel showed, in particular, a plateau for low strain followed by an apparent softening of the material (Fig. 2a). Softening has also been observed and characterized in a recent study by Kalli et al.[22]. One can not exclude the fact that slippage at the plate/sample interface in the rheometer could lead to this apparent softening - we used rough sandpaper during rheological measurement to avoid this effect. We wish to point out that in the case of lower agarose concentration (0.5%), one could reach a critical deformation of about 70% above which macroscopic rupture of the gel is observed, leading to an effective unconfinement of the spheroid (Fig. S1). This rupture could explain the softening observed at high strains. For all 1% agarose samples, deformation remained below this critical 70% strain.

**FIG. 2.**
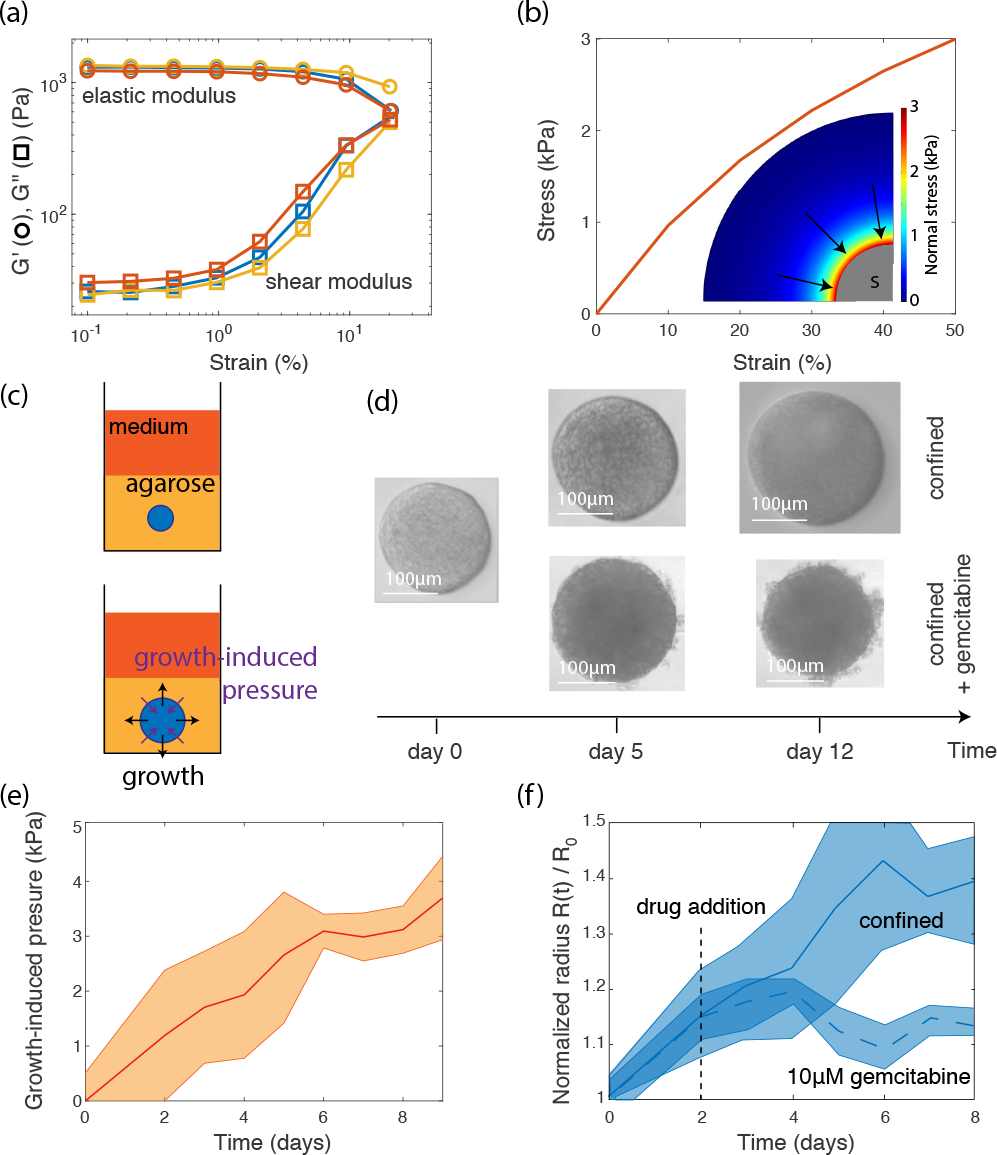
Growth-induced pressure and its impact on gemcitabine efficacy. a. Rheological measurements of elastic modulus G’ (ο) and shear modulus G” (◻) of a 1% low-melt agarose. The different colors correspond to different forces used to hold the sample in position, and show good reproducibility of the data. Softening of the material is observed at large strain. b. Comsol simulation of a spheroid growing in an agarose hydrogel modeled as a Neo-Hookean material with parameters extracted from the rheological measurements. Inset: Heat map of the normal stress onto a spheroid (in grey, S). c. Schematic of an agarose-confined spheroid. d. Representative pictures of a confined spheroid and confined spheroids exposed to gemcitabine. e. Growth-induced pressure as a function of time extracted through the growth of spheroids in 1% low-melt agarose. Median values are plotted normalized to time 0, shading corresponding to ± SD over N ≥ 10 spheroids. f. Quantification of growth of confined spheroids and confined spheroids treated with drug. Median values are plotted, shading corresponding to ± SD over N ≥ 10 spheroids.

Incorporating softening into the Neo-Hookean material properties of a finite-element simulation of a spheroid growing in agarose, as in [22], gave a rough linear increase of normal stress applied onto the spheroid during growth (Fig. 2b). This calibration can be used to extract growth-induced pressure curves from the growth of embedded spheroids (Fig. 2c). Expansion of agarose-embedded spheroids resulted in slower growth (Fig. 2d), similar to what has been described for other embedding solutions[5]. We found that growth-induced pressure rose to the kPa range over a period of a few days (Fig. 2e).

We added 10*μ*M of gemcitabine after 2 days of confined growth and recorded size evolution. Interestingly, we observed that compressed spheroids were less sensitive to chemotherapeutic than freely growing ones with almost no size decrease observed over 5 days (Fig. 2f and Fig. S2): While freely growing spheroids decreased in size by roughly 30-40% (Fig. 1c), we observed less than 10% decrease (Fig. 2f) in size for compressed spheroids. Gemcitabine is a small molecule, slightly smaller than Hoescht 33342 DNA intercalent. We did not not find any difference in Hoescht penetration in between a control and a compressed spheroid (Fig. S3), strongly suggesting that the observed effect of gemcitabine was not due to altered penetration of the drug under compression. Two non-mutually exclusive hypotheses can explain this behavior. The first one entails that compressive stress triggers mechanosensitive pathways directly acting on the effect of the chemotherapeutic, for instance on gemcitabine activation within the cell or import/export rates of the drug[17]. The second hypothesis is that compressive stress would trigger mechanosensors specifically decreasing cell proliferation[9, 21, 23], which would indirectly impact chemotherapeutic efficacy. While capillary western blots could not be performed on agarose-embedded spheroids without biological perturbation, we performed immunostaining of paraffin-embedded samples and observed that compressive stress decreased cell proliferation (Fig. S4).

To the best of our knowledge, there is no study mentioning specific mechanosensitive pathways directly acting on the activity of the drug. Although we cannot rule out their existence, we developed a generic mathematical framework in order to get insight into the potential mechanism limiting drug efficacy under mechanical compression (see Method). We assumed that the total number of cells in the spheroid varied according to cell proliferation and drug-induced death. Cell proliferation was exponentially distributed over the spheroid radius, with a characteristic length *ℓ* (Fig. S4). We observed that cell death induced by the drug is not instantaneous. Indeed cells must be in a proliferative state in order for the drug to be effective. The waiting time for the cell to enter the proliferative phase could be assumed to be exponentially distributed, giving a delay for the drug to be effective after about 24 hours. Assuming that the characteristic proliferation length *ℓ* was smaller than the spheroid radius, we wrote the temporal variation of the spheroid radius *R* as:

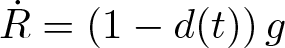

where *g* was the effective growth rate and *d*(*t*) was the death term induced by the drug which included time delay (see Methods for full derivation of the model). The dependence on pressure was only considered for the growth rate *g* which took value *g*_0_ in control conditions (i.e. no compression and no drug). Note that no drug gradients were considered in the model: gemcitabine is equally distributed within the spheroid. We used a single growth curve to fit each parameter individually. In particular, *g* = *g*_0_ was fitted through the control experiment, *g* = *g*_*c*_ was obtained from the spheroid growing under compression, and the drug induced death rate *d* = *d*_0_ was calibrated when the drug was administered to the spheroid in the absence of pressure. With these parameters, we were able to predict, *i.e.* without any fitting parameters, the combined effect of mechanical stress and chemotherapeutic under the assumption of independent effects. The precision of the prediction was scored by a square difference of the measurement with the expected value. Our model remarkably predicted the experimental data (Fig. 3a), with a score χ = 0.94, suggesting that indeed, *d*_0_ did not depend on pressure.

**FIG. 3.**
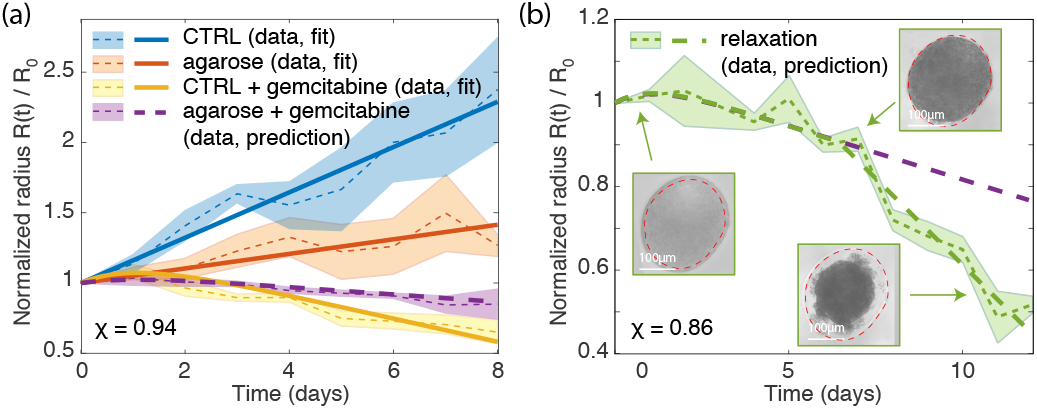
Prediction of the model on confined data. a. Prediction of the model (large dashed purple) on confined spheroids exposed to gemcitabine. The score of the prediction is χ = 0.94. b. Spheroids exposed to gemcitabine can eventually be smaller that the initial inclusion size (pictures in inset). The model (large dashed green) is able to capture the effect of gemcitabine in this case, switching from efficacy of confined cells before 6 days (large dashed purple), to efficacy of unconfined cells after 6 days. χ = 0.86. In both cases, median values are plotted normalized to the time of drug addition, shading corresponding to ± SD over N ≥ 10 spheroids.

Our model predicts that a decrease in compressive stress could lead to an increase in cell proliferation and a higher efficacy of the chemotherapeutic. Although we could not instantaneously relax mechanical stress without any biological perturbation, we took advantage of the fact that treated spheroids decreased in size over time. After 5-6 days of drug treatment, some spheroids saw their radius reducing below the initial inclusion size such that they were not confined anymore and did not experience any compressive stress (pictures inset of Fig. 3b). Note that the spheroids presented in Fig. 3a were always confined. Our model would predict that quiescent cells would re-enter the cell cycle, proliferate faster, and consequently die faster. Our model was perfectly able to capture the experimental data (Fig. 3b, χ = 0.86): a slow initial death rate during compression, followed by a faster one in the unconfined phase.

The excellent agreement of the model with the experimental data is consistent with a mechanism where mechanics would decrease chemotherapeutic efficacy through a modulation of cell proliferation. This mechanism makes two key predictions: If the efficacy of a proliferation-based chemotherapeutic is decreased because of a modulation of cell proliferation, then the observed modulation of efficacy (i) should not depend on the type of drug used, but rather on the fact that the chemotherapeutic targets proliferating cells, and, similarly, (ii) should not depend on the type of mechanical stress applied, but rather on the fact that mechanical stress could decrease cell proliferation. We investigated these two predictions, by treating with a different chemotherapeutic, docetaxel, and applying a different kind of mechanical stress, a mechano-osmotic compression with dextran[8].

Docetaxel is a taxol-based drug which stabilizes microtubules, leading to cell death during M-phase[24]. We confined spheroids in 1% low-melting agarose gel as previously described, and treated the compressed spheroid after 2 days under mechanical compression with 10*μ*M docetaxel. Our model was able to recapitulate the experimental data, the efficacy being reduced for compressed cells, in a predictive manner (χ = 0.94) (Fig. 4a).

**FIG. 4.**
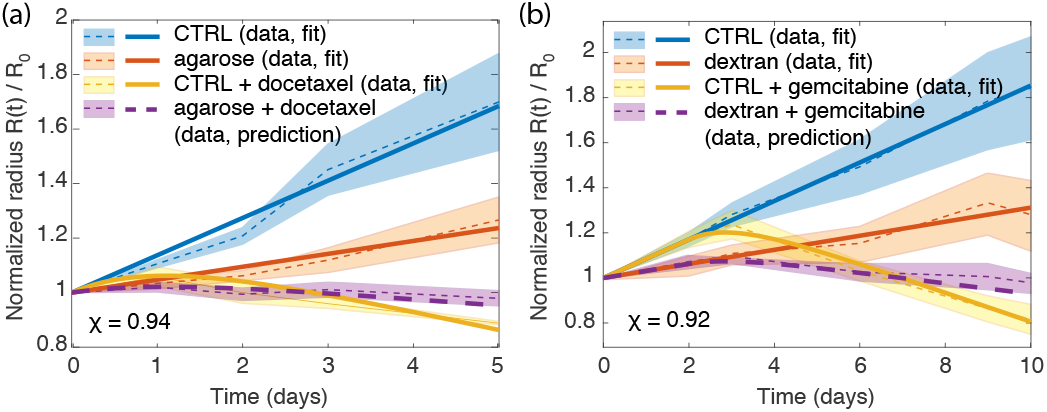
Prediction of the model for (a) confined spheroids exposed to docetaxel (χ = 0.94) and (b) osmotically compressed spheroids exposed to gemcitabine (χ = 0.92). In both cases, median values are plotted normalized to the time of drug addition, shading corresponding to ± SD over N ≥ 10 spheroids.

The addition of high-molecular weight dextran to the culture medium reduced cell proliferation in a similar way as confined growth-induced pressure[8, 23, 25]: We observed that osmotically compressed spheroids restricted cell proliferation to the outermost layers (Fig. S4), and capillary western blots confirmed a decrease in cell proliferation measured by pERK/ERK (Fig. S5). Treatment of osmotically compressed spheroids with gemcitabine showed a qualitatively comparable modulation of efficacy of the drug with growth-induced pressure (Fig. 4b). Once again, our model remarkably predicted the effect of the drug combined to this mechanical stress (χ = 0.92).

Many past studies have identified key features which can eventually lead to drug resistance. Most studies have focused on the inactivation of the drug by the host, the alteration of the drug target, the efflux of the drug from the cell, or DNA alterations that could create *de novo* resistance. All of these mechanisms are cell-centered, but there is an increasing realization that stromal components could also participate in drug resistance.

Our experimental data suggest a novel mechanical-form of drug resistance which could arise from tumor-stroma mechanical interaction. Triggering of signaling cascade reducing the activity of a given chemotherapeutic under mechanical stress seems not to be needed for resisting the drug. Rather, the efficacy of a drug can be directly altered by a mechanical-control of cell proliferation which can occur through dedicated sensors. The note-worthy theoretical prediction of the experimental data, with an underlying assumption that the effects of mechanical stress and drug activity are uncoupled, strongly supports this mechanism[26]. We observe that growth-induced pressure, a highly common type of mechanical stress present in most solid tumors, can modulate the efficacy of chemotherapeutics acting on different parts of the cell cycle. Moreover, an osmotic compression, stress of very different origin and sensing which could arise *in vivo* due to accumulation of oncotic pressure[27], also leads to similar modulation of chemotherapeutic. There are now evidences that quiescence is one major mechanism leading to drug resistance[28] and that mechanics can turn cells towards quiescence [8, 29]. However, to our best knowledge, no quantitative models linking mechanics to drug resistance, through a direct modulation of proliferation, have been proposed so far. Our data can potentially explain the reduced effect of chemotherapeutics of mechanically-stressed cells[13, 14].

Most if not all solid tumors experience compressive stress. While this stress can be heterogeneous *in vivo*, some cells could be compressed and create pockets of drug resistance inside a tumor. Within the framework of a mechanical-form of drug resistance, it appears clear that the mechanical modulation of the microenvironment could be an interesting therapeutic option, although much work remains to be done. For instance, hyaluronidase, which is currently under clinical trial, may have a very different effect than only modulating tumor perfusion: by reducing matrix swelling and subsequent compressive stress experienced by cells, it could also modulate cell proliferation. A direct mechanical-modulation of drug efficacy through cell proliferation could appear particularly deleterious, notably because it does not rely on any specific gene alteration targeting the mode of action of the drug, and should be carefully accounted for during cancer treatment. This mechanism calls for a better understanding of the mechanosensors at play during proliferation reduction under mechanical stress: A therapy targeting these sensors to enforce cell proliferation under mechanical stress coupled with a proliferation-driven chemotherapeutic could represent an appealing strategy to battle compressed tumors.

## Acknowledgement

The authors would like to thank Davide Ambrosi for useful discussions. IR, SA, JGG and MD performed biological experiments; AP, CB, CR and HDA performed rheological experiments and simulations; PM, HH and MD did the theoretical modeling; and IR, PM, HDA, HH, JGG and MD analyzed the data and wrote the article. H.H. and P.M. acknowledge the funding support of MicMode-I2T (01ZX1710B) by the Federal Ministry of Education and Research (BMBF). JGG and MD thank the Fondation Toulouse Cancer Santé, Plan Cancer Inserm, and Cancéropole Grand Sud Ouest for financial support.

