## Supplementary information for "Mechanical-control of cell proliferation increases resistance to chemotherapeutic agents"

[6] *Laboratoire d'Excellence TouCAN*

[7] *Univ Lyon, Univ Claude Bernard Lyon 1, CNRS, Institut Lumière Matière, F-69622, Villeurbanne, France*

[†] *The authors contributed equally*

[*] *To whom correspondance should be sent:*

**Cell culture, spheroid formation.**

A338 cell line, derived from a murine pancreatic tumor with an activating mutation of KRas oncogene ($KRas^{G12D}$), were culture in DMEM (Sigma-Aldrich), supplemented with 10% FBS (Sigma-Aldrich), at 37°C and 5% $C0_2$. Spheroids were formed following a classical agarose cushion protocol[1]. Briefly, 200$\mu$L of 1% agarose was polymerized in each wells of a 48-well plate. 600$\mu$L of a cell suspension (typically 1800 cells/mL) was then added in each well. After 2 days, spheroids formed at the bottom of each well and had a typical size of about 200$\mu$m.

**Embedding in agarose hydrogel.**

A 48-well plate was placed on ice. We prepared a low-melting agarose solution of 2% concentration and left it at 37° to thermalize. We removed 400$\mu$L of medium from a well of a 48-well plate containing spheroids of 2/3 days old. The remaining 200$\mu$L of medium containing the spheroid was then mixed with 2% low-melting agarose within the pipette. The 400$\mu$L solution was placed on the 48-well plate on ice, to allow for rapid polymerization of agarose at a final concentration of 1%. We found that this step was necessary to obtain a fully-embedded spheroid: if the polymerization occured at room temperature, the spheroid sedimented most of the time at the bottom of the well, and was not embedded in 3D. Note that we tried to polymerize low-melting agarose directly on top of a spheroid which was on the agarose cushion, but found similar result with a spheroid that sedimented: the mechanical stress around the spheroid was not homogeneous. The control spheroids were subjected to the same manipulation. Importantly, we compared the size the of the spheroid before embedding and right after manipulation and being on ice and found no significant different in its size.

**Application of dextran osmotic stress.**

After 2 days of spheroid formation, 300$\mu$L of culture medium was replaced by 300$\mu$L of 70kDa dextran at 40g/L concentration to reach a final concentration of 20g/L, corresponding to a exerted osmotic stress of 1kPa[1].

**Drug treatment.**

After 2-3 days of growth under mechanical stress, either coming from agarose confinement or osmotically from dextran, drug was added to the culture medium. The vehicle of the drug was added to the control. Gemcitabine (IUCT-O, Toulouse) was added at a final concentration of 10$\mu$M, and docetaxel (IUCT-O, Toulouse) was added at a final concentration of 10$\mu$M.

**Agarose sample preparation for rheological measurements.**

A solution of low-melting agarose (A3414, Sigma-Aldrich) diluted in distilled water was prepared by autoclaving the sample at 120°C (1% (w/v)) for 15 min. Agarose pads were then molded in 35x10 mm Petri dish (353001 Falcon) and left at room temperature for a few minutes. A glass coverslip and a weight were placed on the top of the Petri dish to obtain a perfect flat pad after unmolding. The sample was left to polymerize in the fridge overnight before rheological measurements.

**Rheological characterization of agarose.**

The viscoelastic properties were determined using oscillatory deformation applied by a stress-controlled rotational rheometer (Anton Paar MCR 301). The measurements were performed in a plate-plate (PP) geometry at room temperature. Upper and lowerplate diameters respectively of 42mm and 64mm were used. The dimension of the gap was in the range of 8.5-9 mm (a little smaller than the dimension the agarose sample). Sandpaper with a roughness of 40$\mu$m was glued to the plates to avoid slippage of the sample upon shearing. The normal force associated with the confinement varied between 0.05 to 1.35 N. We performed amplitude sweep tests at a frequency of 1Hz, the duration of the measurement was automatically set by the rheometer. We applied a correction on the shear modulus given by the software, as our sample was smaller than the plate used:

$$G'_{\text{real}} = \left(\frac{R}{r}\right)^4 G'_{\text{software}} \tag{1}$$

R being the radius of the measurement plate (21 mm) and r being the radius of our sample (17.5mm). This gives:

$$G'_{\text{real}} \sim 2G'_{\text{software}} \tag{2}$$

**Comsol simulations.**

Finite element simulations were done using Comsol multiphysics with the solid mechanics module in a cylindrical geometry. We used a Neo-Hookean material for the agarose gel with a shear modulus value taken from the rheology experiment (2800 Pa). The bulk modulus was taken as $10^6$ Pa, and the material was considered as nearly incompressible. The deformation is imposed at the inner radius, and null stress is imposed on the exterior surface of the gel (see inset of Fig2b. for a scheme of the geometry).

**Testing drug penetration.**

Spheroids were embedded in agarose hydrogels as mentioned above. After 2 days of confined growth, corresponding to the time point of drug addition, we added Hoescht-33342 (Sigma), a methylated form of DAPI which freely enters the cell and the nucleus. We performed confocal slicing using a 100mW laser and a Yokogawa psinning disk on a Leica microscope, and selected the optical plane in the middle of a spheroid (Fig. S3). We observe a similar penetration of Hoescht inside the control and compressed spheroids, suggesting no decrease in perfusion at this size-scale.

**Biological characterization of samples.**

We performed two types of biological characterization of the samples: capillary western blots, when the spheroids could easily be taken from the medium, and immunostaining of sections of spheroids.

Capillary western blot having an increased sensitivity enables to use less biological sample per read. Typically, 5 spheroids were used to get enough signal. At the end of the experiment the spheroids to analyzed were lysed. 400$\mu$L of medium was removed from every well, then the spheroids from the same experimental condition were gathered together in a Falcon tube, spun down by soft centrifugation (200 g) and washed with cold PBS three times (5 minutes each). It was important to remove most of the liquid from the spheroids: the excess liquid would otherwise dilute the lysis buffer in the next steps. At this point, the dried samples could be frozen at -80°C or lysed directly. A lysis buffer was prepared with 150mM NaCl, 50 mM Tris, 1mM EDTA, 1% Triton; 2 mM of Dithiothreitol (DTT), 2mM of NaF (Serine/Threonine phosphatase inhibitor), Sodium Orthovanadate (Tyrosine Phosphatase inhibitor) 4mM and a tablet of complete Roche proteases inhibitor cocktail were added just before lysing. Spheroids were then incubated on ice with 200$\mu$L of lysis buffer and vortexed regularly for 15 minutes. Then, the samples were centrifuged 5 minutes at 9500 g - 4°C and the supernatant was transferred to an Eppendorf tube. The machine Automated Western Blot from Protein Simple Company (Wes) and its corresponding preparation kits were used for a high performance capillary western blot. In Wes, proteins were size separated in capillaries, where they were incubated with primary and (HRP-conjugated) secondary antibodies and finally with Luminol/peroxidase. The produced chemiluminescence was detected at multiple exposure times and automatically quantified by the Compass software. The samples were appropriately diluted with 0.1 X sample buffer. Then, 1 part 5X Fluorescent Master Mix with 4 parts diluted lysate in a microcentrifuge tube (final concentration 0.4 mg/mL for chemiluminescence) was combined producing enough diluted sample volume required for assay. The samples were denatured in a specific hot plate at 95oC for 5 minutes. ERK (Extracellular signal Regulated Kinase) was used as proliferation marker (Cell Signaling), cyclin dependent kinase inhibitor 1B (p27) was used to identify proliferation inhibition (Sant Cruz) and cleaved-caspase 3 was used as an apoptosis marker (Cell Signaling). The loading quantity was controlled by the total volume of analyzed spheroids. The antibodies were diluted 1:50 with the antibodies buffer (Protein Simple company). The solution volume was chosen based on the samples number to be analyzed in the assay. Finally, we prepared the solution luminol-peroxide with the ratio 1:1. Protein samples, blocking reagent, washing buffer, primary antibodies, secondary antibodies, and chemiluminescent substrate were dispensed into the assay plate according to the manufacturer's protocol. Assay plate was then loaded into the instrument, and proteins were separated into individual capillaries. Protein separation and detection was performed automatically on the individual capillaries. The data were analyzed by Compass software.

Whole spheroids could be analyzed by immunostaining. Paraffin-embedding was preferred over cryo-embedding for ease of sectioning. Free or agarose-embedded spheroids were embedded in 1% low melt agarose, then fixed in 10% neutral-buffered formalin (SIGMA) o/N, stored in Ethanol 70% then embedded in paraffin. Next they were serially sectioned (4 $\mu$m), and, on the sections with the largest diameter, immunostainings were conducted using standard methods on formalin-fixed, paraffin-embedded tissues. Antigen retrieval was done with sodium citrate buffer and antibody dilution were carried out (Ki67, Abcam 1:200; cleaved caspase 3, Cell signaling 1:50). All rabbit primary antibodies were revealed using a Cell Signaling Signal Boost system followed by AEC incubation (Dako). For IHC images a transmitted light microscope was used.

**Mathematical model to predict the combined effect of compressive stress and chemotherapeutic.**

We assumes that the total number of cells $N$ varies in the spheroid according to

$$\dot{N} = \dot{N}_p - \dot{N}_d - \dot{N}_a, \tag{3}$$

where $\dot{N}_p, \dot{N}_d, \dot{N}_a$ are the variation of cell proliferation, death due to drug action and apoptosis, respectively. We assumes that these terms had the following functional form:

$$\dot{N}_p = \gamma \int_V \rho\omega \, dV, \quad \dot{N}_d = \delta \int_V \rho\omega \, dV, \quad \dot{N}_a = \alpha \int_V \rho \, dV, \tag{4}$$

where $\rho$ was the cellular density, $\gamma, \delta, \alpha$, were the proliferation, drug-induced death and apoptosis rates, respectively, and $\omega$ was the radial distribution of proliferating cells in the spheroid (see the SI of [2]). In the following we assumed that cell proliferation was exponentially distributed over the spheroid radius, with a characteristic length $\ell$:

$$\omega = \exp\left(-\frac{R-r}{\ell}\right), \tag{5}$$

where $r$ is the spheroid radial coordinate. We enforced spherical symmetry and write

$$\dot{N} = \frac{4}{3}\pi\dot{\rho}R^3 + 4\pi\rho R^2\dot{R}, \tag{6}$$

and

$$\dot{N}_p - \dot{N}_d = 4\pi\rho(\gamma-\delta)\int_0^R e^{-\frac{R-r}{\ell}} r^2 \, dR, \quad \dot{N}_a = 4\pi\rho\alpha\int_0^R r^2 \, dR. \tag{7}$$

By calculating the integrals we obtain

$$\frac{\dot{\rho}}{\rho}\frac{R}{3} + \dot{R} = (\gamma-\delta)\ell\left[1 - 2\frac{\ell}{R} + 2\frac{\ell^2}{R^2}\left(1 - e^{-\frac{R}{\ell}}\right)\right] - \frac{\alpha}{3}R. \tag{8}$$

We assumed that the characteristic proliferation length was small if compared to the radius. Under this condition, equation in (8) simplified to

$$\frac{\dot{\rho}}{\rho}\frac{R}{3} + \dot{R} = (\gamma-\delta)\ell - \frac{\alpha}{3}R, \quad \ell \ll R. \tag{9}$$

By neglecting variations in cell density, equation (9) reduced to a classical von Bertalanffy growth equation[3, 4]. The variation of cell density $\dot{\rho}$ was linked to the pressure that was exerted on the spheroid. When spheroids were compressed by Dextran, this term can be set to zero over the timescale of spheroid growth. In the case of compression by agarose, however, pressure changed over time needed to be taken into account. This happened because the spheroid deformed the agarose while growing and the gel responded elastically by exerting a pressure on the spheroid. We assumed that under the deformation regime that occured in our experiments the agarose can be described by a linear elastic behavior. Therefore, the variation of density $\dot{\rho}$ and pressure $\dot{P}$ could be described by the following relation[5, 6]

$$\dot{\rho} = \frac{\rho}{\chi}\dot{P} = \frac{\rho}{\chi}\frac{4}{3}\frac{E}{R}\dot{R}, \tag{10}$$

where $\chi$ was the spheroid compression bulk modulus. Equation (9) became

$$\left(1 + \frac{4}{9}\frac{E}{\chi}\right)\dot{R} = (\gamma - \delta)\ell - \frac{\alpha}{3}R, \tag{11}$$

$$\dot{R} = \frac{\gamma - \delta}{\beta}\ell - \frac{\alpha}{3\beta}R, \tag{12}$$

$$\dot{R} = \left(\frac{\gamma}{\beta}\ell - -\frac{\alpha}{3\beta}R\right) - \frac{\delta}{\beta}\ell, \tag{13}$$

where $\beta = 1 + 4E/(9\chi)$. Note that the proliferation rate and characteristic length $\gamma$ and $\ell$, as well as the apoptotic rate $\alpha$, depended on pressure and should be written as a function of spheroid radius in a way similar to what we did for equation (10). However, previous experimental work[2, 5] suggested that the latter quantities reached an asymptotic value after a few kPa of pressure. Similar pressure values were obtained in the agarose gels that we considered even for small deformations, so that we could neglect the dependence of $\gamma, \ell$ and $\alpha$ on the spheroid radius and use directly their asymptotic values.

Since our experiments were performed over a timescale and size-scale in which apoptotic death in the spheroid was not thoroughly developed, i.e. $\alpha t/(3\beta) \ll 1$, we rewrote equation (13) as

$$\dot{R} = (1 - d)g, \tag{14}$$

where $g$ was the effective net growth rate (including the early effects of cell apoptosis in the spheroid), and $d$ was a death term induced by the drug. In writing (14) we assumed that $d$ was independent of the external pressure acting on the spheroid. The dependence on pressure was only considered for the growth rate $g$, which took the value $g_0$ in control conditions (i.e. no compression and no drug) and $g_c$ in the compressed case. We prescribed the death term induced by the drug $d$ to depend on time as

$$d = \begin{cases} 0, & \text{for } t < t_0 \\ d_0[1 - e^{-r(t-t_0)}], & \text{for } t \geq t_0, \end{cases} \tag{15}$$

where $t_0$ is the time at which the drug was administered and $r$ was a standard proliferation rate set to $r = 1\text{d}^{-1}$. Indeed, gemcitabine and docetaxel were effective on cells that had entered the S-phase or M-phase, respectively, of the cell cycle. By choosing $d$ according to (15) we assumed that the probability of entering the S-phase followed an exponential distribution, with a rate equal to the proliferation rate. Finally, to quantify the quality of the model predictions we introduced the score

$$\chi = 1 - \sqrt{\sum_i (R_i^{\text{e}} - R_i^{\text{m}})^2}, \tag{16}$$

where the summation was performed on the experimental points of the treatment curve in the presence of compression and $R^{\text{e}}$ and $R^{\text{m}}$ were the radii from the experiment and the model, respectively.

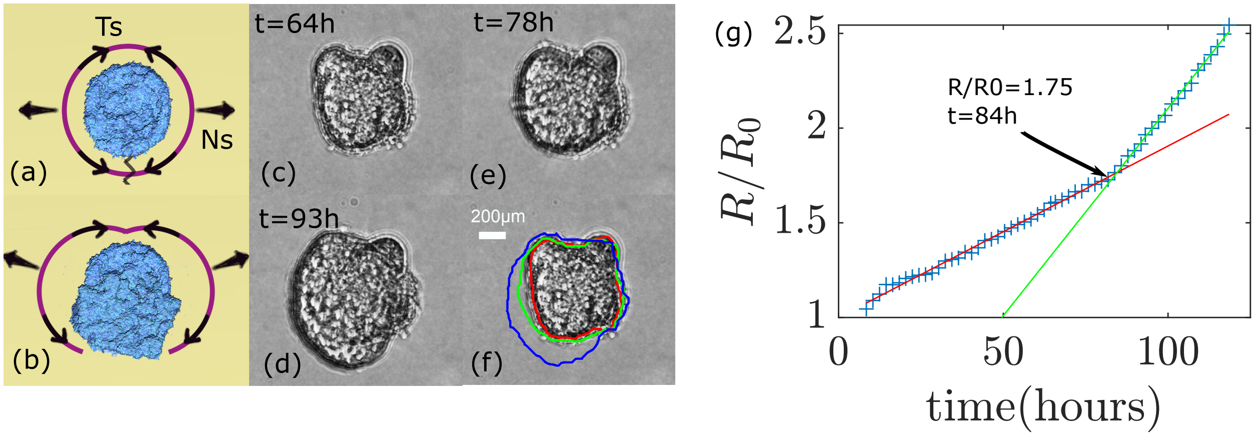


Supplementary figure S 1. a. As aggregates grew in agar gel, they induced a tensile and normal stress (Ts and Ns), that could eventually lead to gel rupture (b). c-f. Example of such a rupture in 0.5% agar gel. f. Superimposition of the aggregate's contour of image (c-d-e) respectively in red green and blue. Note that the rupture was not optically visible. Its signature resided in the asymmetric growth of the aggregate after its occurence (growth in the down left corner here) and in a rupture of the growing slope. g. Normalized radius of the aggregate as a function of time, a clear rupture in the growth slope is seen at time t=84h, certainly corresponding to the appearance of a fracture in the gel.


[1] F. Montel, M. Delarue, J. Elgeti, L. Malaquin, M. Basan, T. Risler, B. Cabane, D. Vignjevic, J. Prost, G. Cappello, *et al.*, Physical review letters **107**, 188102 (2011).
[2] F. Montel, M. Delarue, J. Elgeti, D. Vignjevic, G. Cappello, and J. Prost, New Journal of Physics **14**, 055008 (2012).
[3] L. Von Bertalanffy, The quarterly review of biology **32**, 217 (1957).
[4] M. Marušić, Mathematical Communications **1**, 175 (1996).
[5] K. Alessandri, B. R. Sarangi, V. V. Gurchenkov, B. Sinha, T. R. Kießling, L. Fetler, F. Rico, S. Scheuring, C. Lamaze, A. Simon, *et al.*, Proceedings of the National Academy of Sciences **110**, 14843 (2013).
[6] L. Landau and E. Lifshitz, "Theory of elasticity. butterworth," (1986).
[7] M. Delarue, F. Montel, D. Vignjevic, J. Prost, J.-F. Joanny, and G. Cappello, Biophysical journal **107**, 1821 (2014).

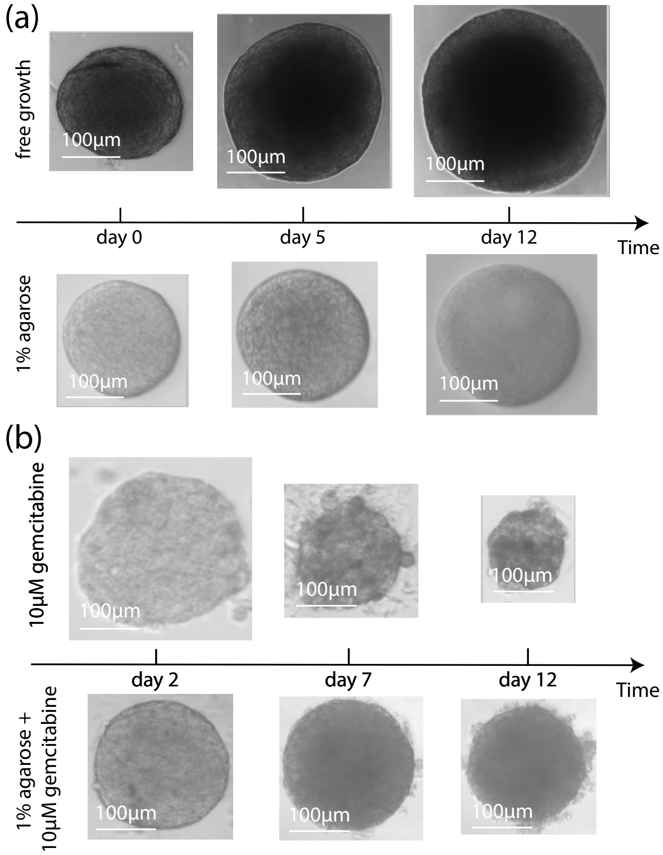


Supplementary figure S 2. Representative pictures of spheroid growing over time. a. Controlled and confined growth in 1% low-melting agarose. b. Spheroids treated with 10$\mu$M gemcitabine with or without confinement.

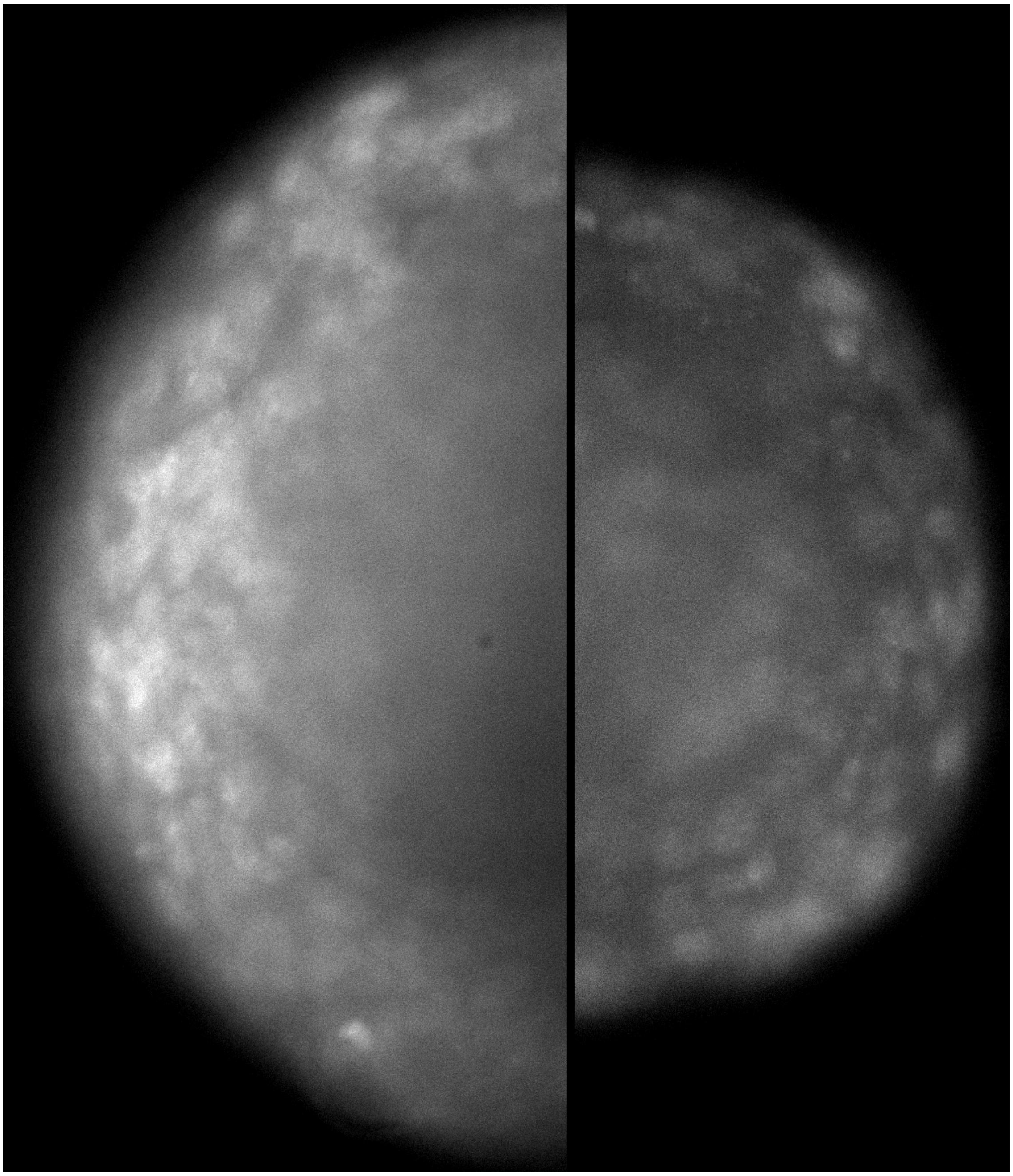

Supplementary figure S 3. Confocal slice inside a control and a mechanically compressed spheroid in DAPI channel. Hoescht-33342 (Sigma) was added to the culture medium 1h before imaging. The section represents the middle of the spheroids. We can observe in both spheroids that Hoescht, bigger than gemcitabine, was able to penetrate the spheroids and stain the nucleii of the first 4-5 layers, showing no decrease in penetration at this size-scale. Beyond this limit, it is hard to conclude on Hoescht penetration due to imaging limitations.

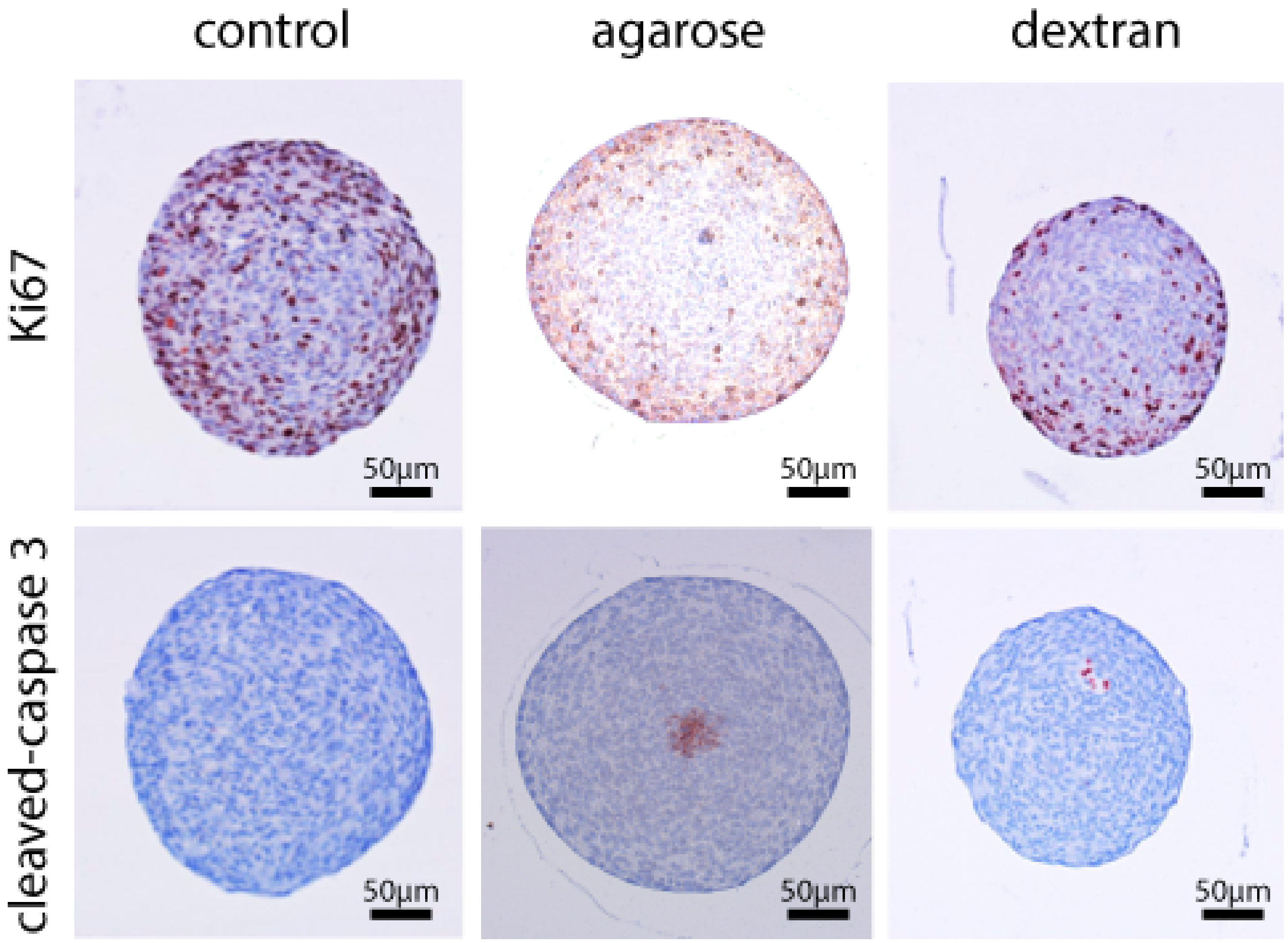


Supplementary figure S 4. Representative immunostaining of sections from paraffin-embedded spheroids. A control, agarose-embedded, and osmotically stressed spheroid was stained with Ki67 for proliferative cells[1], and with cleaved-caspase 3 for cell death. Note that the agarose-embedded was a bit lighter than the control or dextran: the agarose slightly interfered with the staining. Pictures showed that compressive stress, either mechanical or osmotic, decrease cell proliferation, without a significant increase in cell death. In particular, no cell death was observed for the control, suggesting that no strong nutrient or oxygen gradients was present for this size of spheroid.

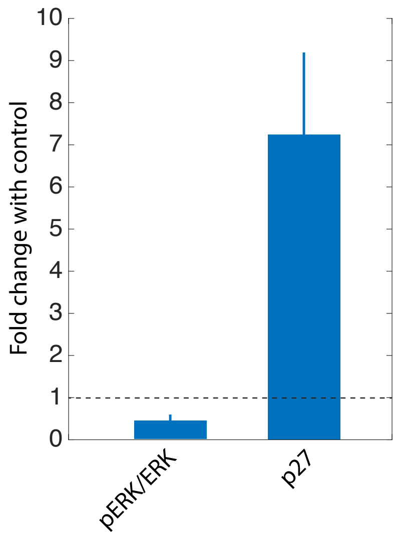


Supplementary figure S 5. Capillary western blots were performed on spheroids subjected to 1kPa of dextran osmotic stress. Phosphorylated-form of ERK over total ERK was used as a proliferation marker, and p27, a CDK inhibitor, as a proxy for mechanical-induction of quiescence [7]. The data were normalized to the control condition, without dextran. We observed a decrease in the amount of proliferative cells, consistent with the immunostaining of the previous figure, observed by a decrease in the amount of pERK/ERK and an increase in the amount p27.
